# Network component analysis reveals developmental trajectories of structural connectivity and specific alterations in autism spectrum disorder

**DOI:** 10.1101/100164

**Authors:** Gareth Ball, Richard Beare, Marc L. Seal

## Abstract

The structural organisation of the brain can be characterised as a hierarchical ensemble of segregated modules linked by densely interconnected hub regions that facilitate distributed functional interactions. Disturbances to this network may be an important marker of abnormal development. Recently, several neurodevelopmental disorders, including autism spectrum disorder (ASD), have been framed as disorders of connectivity but the full nature and timing of these disturbances remain unclear.

In this study, we use non-negative matrix factorisation, a data-driven, multivariate approach, to model the structural network architecture of the brain as a set of superposed subnetworks, or network components.

In an openly available dataset of 196 subjects scanned between 5 to 85 years we identify a set of robust and reliable subnetworks that develop in tandem with age and reflect both anatomically local and long-range, network hub connections. In a second experiment, we compare network components in a cohort of 51 high-functioning ASD adolescents to a group of age-matched controls. We identify a specific subnetwork representing an increase in local connection strength in the cingulate cortex in ASD (t=3.44, p<0.001).

This work highlights possible long-term implications of alterations to the developmental trajectories of specific cortical subnetworks.

## Introduction

Diffusion MRI allows the non-invasive inference of white matter pathways in the human brain. At a millimetre-scale, the structural connections between brain regions can be conceptualised as a complex network and interrogated with graph theoretical approaches (Bullmore and Sporns 2009). This has led to the broad characterisation of the macroscale organisation of the mammalian brain as a near-decomposable system built on multiple, parallel and partially segregated modules (Simon 1962; Meunier et al. 2010). Network modules are organised hierarchically (Bassett et al. 2011) and linked by a set of overarching, densely interconnected hub regions that facilitate distributed interactions across the network (Sporns et al. 2005; van den Heuvel and Sporns 2011; Bullmore and Sporns 2012).

This view is supported by evidence that cerebral regions can be clustered together based on the extent of their shared connections into communities, or modules (Hilgetag et al. 2000; Bullmore and Sporns 2012). Anatomical connectivity between regions reflects a shared functional specialisation (Hilgetag et al. 2000; Honey et al. 2009), and connected regions tend to have similar metabolic demands, and gene expression profiles (Vaishnavi et al. 2010; French and Pavlidis 2011; Collin et al. 2013; Fulcher and Fornito 2016). Furthermore, anatomically connected regions tend to mature in tandem across development (Raznahan et al. 2011), resulting in common patterns of cortical growth and functional coordination over the lifespan (Hagmann et al. 2010; Zielinski et al. 2010; Alexander-Bloch et al. 2013). Taken together, this evidence suggests that connections within complex brain networks can be decomposed, or clustered, into subnetworks that link modules with distinct roles and developmental trajectories.

Long-distance cortico-cortical connections are established during gestation, and complex network architecture is evident at birth (Ball et al. 2014). The effects of cerebral maturation on increasingly distributed connectivity is marked in the first years of life (Yap et al. 2011), after which the large-scale topological organisation of the structural connectome remains relatively stable (Dennis et al. 2013; Baker et al. 2015). Over the full lifespan, measures of network efficiency and modularity follow a distinct inverted U trajectory, peaking in the third decade and mirrored by microstructural markers of the underlying white matter (Imperati et al. 2011; Kochunov et al. 2012; Zhao et al. 2015). In elderly individuals, although network topology remains relatively consistent with younger adults, simulations suggest a preference for local communication compared to long-range hub-to-hub connectivity, correspondent to evidence from functional analyses (Cao et al. 2014; Perry et al. 2015).

The early establishment of structural connectivity and long-term stability of structural networks suggests that disturbances to network organisation may be an important marker of abnormal cerebral development. A number of neurodevelopmental disorders, including autism spectrum disorder (ASD) and attention deficit hyperactivity disorder (ADHD), have been linked to alterations in the development of structural and functional brain connectivity (Konrad and Eickhoff 2010; Tomasi and Volkow 2012; Ecker et al. 2015). ASD is a complex, multifactorial disorder characterised by social, behavioural and language impairments evident from an early age. Although the aetiology of ASD remains unknown, neuropathological studies have identified cortical alterations including laminar and columnar disorganisation and increased neuronal density in frontal, temporal and cingulate cortices (Bailey et al. 1998; Casanova et al. 2002; Stoner et al. 2014; Uppal et al. 2014). Early evidence from MRI studies suggested that head growth is accelerated in ASD during infancy but differences appear to dissipate with age (Courchesne et al. 2001; Ecker et al. 2015). More recently, ASD has been framed as a disorder of connectivity (Belmonte et al. 2004; Vissers et al. 2012), based on accumulating evidence of disruptions to both functional and structural networks in autistic populations (Rudie et al. 2012; Mueller et al. 2013; Supekar et al. 2013; Di Martino et al. 2014). Although the nature and extent of these alterations remain unclear with a number of conflicting observations, previous studies have described complex patterns of disrupted white matter organisation in ASD that appear to be dependent on age and mirrored by evidence of both hypo -and hyper-connectivity between functional networks and differences in electrophysiological recordings (for review, see Vissers et al. 2012).

In this study, we apply an unsupervised and data-driven approach to model complex networks derived from whole-brain tractography as a set of components, or subnetworks, that vary together across the population. We first demonstrate that network components can be robustly and reliably identified in a large cohort, before exploring the developmental trajectories of each component across the human lifespan. In a second experiment we test the hypothesis that ASD is associated with increased structural connectivity. By identifying a set of network components in a group of adolescents with high-functioning ASD and age-matched, typically-developing controls, we find a specific cortical subnetwork with significantly increased connection strength in the autistic population.

## Methods

### Data

Preprocessed connectivity data were downloaded from the USC Multimodal Connectivity database (http://umcd.humanconnectomeproject.org) (Brown et al. 2012). Full MRI acquisition and image processing details are given elsewhere (Brown et al. 2012; Rudie et al. 2012) but are reported in brief below.

#### NKI-Rockland lifespan sample

In total, connectivity matrices from 196 healthy participants (114 male; age range: 4-85y) were available, collected as part of the NKI/Rockland lifespan study (Nooner et al. 2012). Diffusion MRI was acquired at 3 T with 64 gradient directions and the following parameters: TR, 10000 ms; TE, 91 ms; voxel size, 2 mm^3^; *b*-value, 1000 s/mm^2^. After correction for motion and eddy current distortions using linear registration, diffusion tensors were modelled and tractography performed using fibre assignment by continuous tracking (FACT) with an angular threshold of 45° (Mori et al. 1999).

To construct each connectivity matrix, 188 regions-of-interest (ROI) were defined using a group-based functional MRI (fMRI) parcellation (Craddock et al. 2012), and structural connectivity was calculated as the total number of streamlines connecting any two ROI (Brown et al. 2012). Prior to analysis, ROI in the brainstem were removed and fibre counts were log-transformed resulting in a 182 × 182 connectivity matrix for each participant.

#### UCLA autism sample

Connectivity data from a total of 43 typically developing children and adolescents (36 male; age range: 8.9-17.9y) and 51 with high-functioning ASD (45 male; age range: 8.4–18.2 y) were available for analysis (Rudie et al. 2012). 3 T DTI data were acquired with: 32 gradient directions; TR, 9500 ms; TE, 87 ms; voxel size, 2 mm^3^; *b*-value, 1000 s/mm^2^. Motion and eddy current correction, diffusion tensor modelling and tractography were performed as above but with an angular threshold of 50°.

ROI were defined as 10 mm radius spheres placed at 264 coordinates in MNI space and transformed to individual diffusion data (Power et al. 2011) and connectivity was defined as streamline count between connected ROI. As above, fibre counts were log-transformed before analysis, resulting in a 264 × 264 connectivity matrix for each participant.

### Non-negative matrix factorisation

Non-negative matrix factorisation (NMF) is an unsupervised, multivariate decomposition technique that models an *n* × *m* data matrix, *V*, as the product of two non-negative matrices: *W* and *H*:

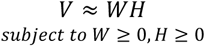

where *n* is the number of features and *m* is the number of samples and *W* and *H* have the dimensions *n* × *r* and *r* × *m* respectively, where *r* is the number of network components or basis images (Figure 1). The optimal solution is sought by iteratively updating *W* and *H* to minimise the (Euclidean) distance between the original and reconstructed matrices, subject to non-negativity constraints:

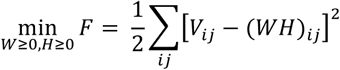

**Figure 1:**
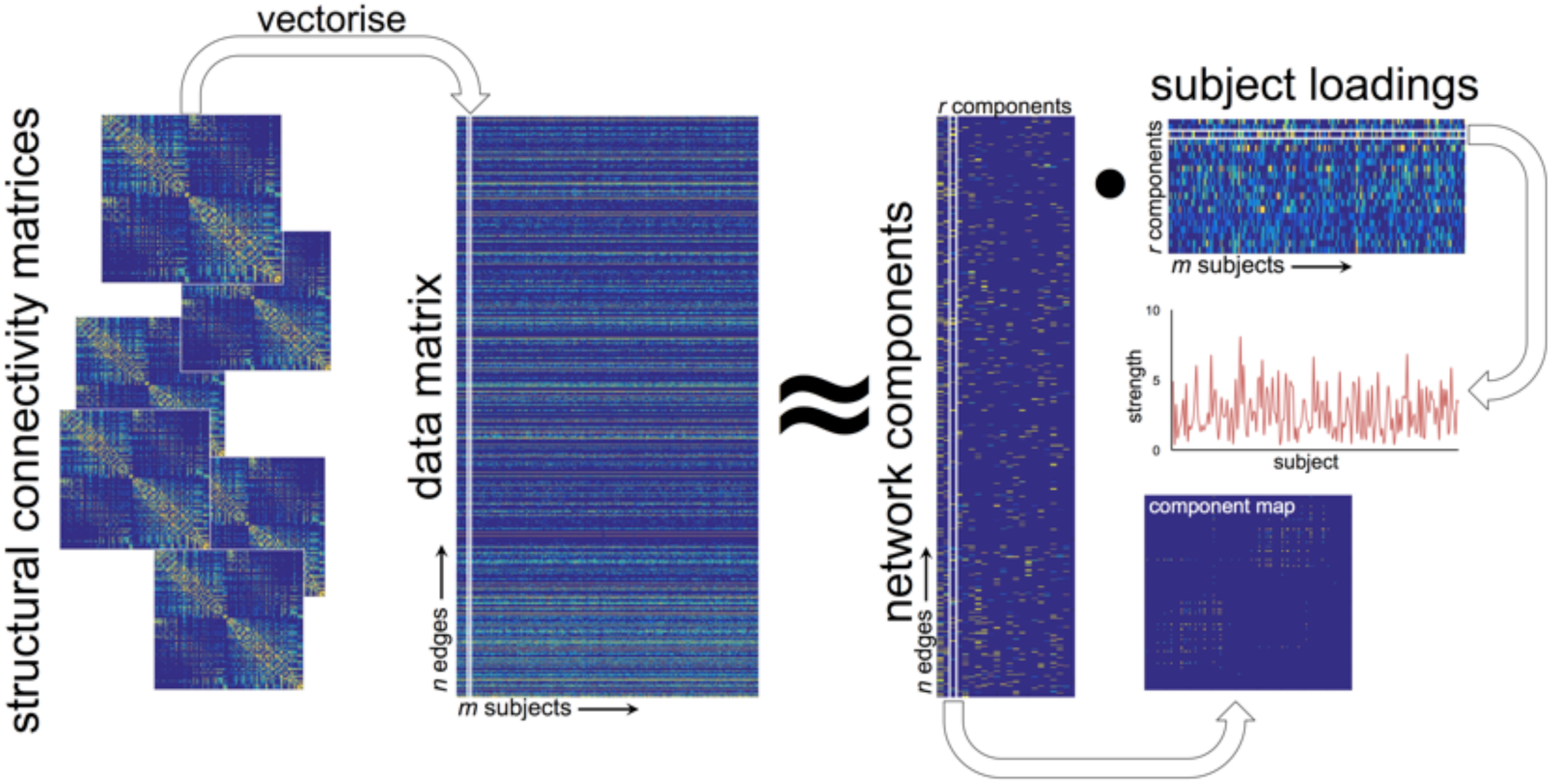
Projective NMF pipeline. Individual connectivity matrices are concatenated into a large data matrix. Projective NMF is used to decompose the data into a set of network components. A map of connections shows the topological organisation of each component, or subnetwork, and a subject-specific weighting estimates the component’s contribution each individual’s full network.

Generally, *r* < min(*m, n*), thus *WH* represents a low-rank approximation of the original data in *V* (Lee and Seung 1999). NMF offers a natural setting for exploration of data that is inherently non-negative and, as such, is particularly well–suited to neuroimage analysis allowing an intuitive understanding of image-derived, non-negative features including e.g.: tissue volume, image intensity, cortical thickness, fractional anisotropy and structural connectivity (Sotiras et al. 2015).

In a recently introduced variant, projective NMF (PNMF), the subject loading matrix, *H* is replaced with *W^T^ V* such that:

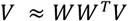

PNMF confers a number of benefits over standard NMF including fewer learned parameters, and increased sparsity and orthogonality of the resulting component matrix *W* (Yang and Oja 2010).

In terms of a network analysis, PNMF results in a set of highly orthogonal network components, each comprising a sparse set of (topologically) localised connections (i.e.: the edge structure of different components does not overlap) and a corresponding subject-specific weighting (Figure 1). Together, these elements can then be combined to approximately reconstruct the full connectivity network of any given subject.

In addition, Yang et al. introduced a method to estimate the rank of the factorising matrix, *W*, using automatic relevance determination (ARD-PNMF; Yang et al. 2010). Here, we employ ARD-PNMF to perform an exploratory analysis of network structure and extract a set of effective network components for further analysis.

### Network decomposition

The analysis pipeline is shown in Figure 1. For each study, structural connectivity networks were vectorised and collated into an *n × m* matrix before normalising to [0,1]. To reduce computation time and remove noisy connections, edges that were present in less than 10% of the study population were removed. ARD-PNMF was initialised with non-negative double singular value decomposition, a procedure that speeds up NMF convergence compared to a random initialisation and ensures consistent results across runs (Boutsidis and Gallopoulos 2008). We chose an initial rank estimate of 50 and performed a maximum of 20,000 PNMF iterations, or until the algorithm converged.

Network decomposition was performed in Matlab R2015b using PNMF code available at: sites.google.com/site/zhirongyangcs/pnmf and NNDSVD code available at: www.boutsidis.org/software.html.

### Simulations

To demonstrate the application of PNMF to structural connectivity data, we performed a set of simulation experiments. We created a set of 150 ‘networks’, each comprising a weighted combination of six network components (Figure 2). Each network component was constructed by adding binary edges between 10-20 randomly selected nodes in a 100 × 100 empty network. The weighted contribution of each component to an individual network was varied according to a set of predefined patterns that varied across the population (Figure 2A).

**Figure 2:**
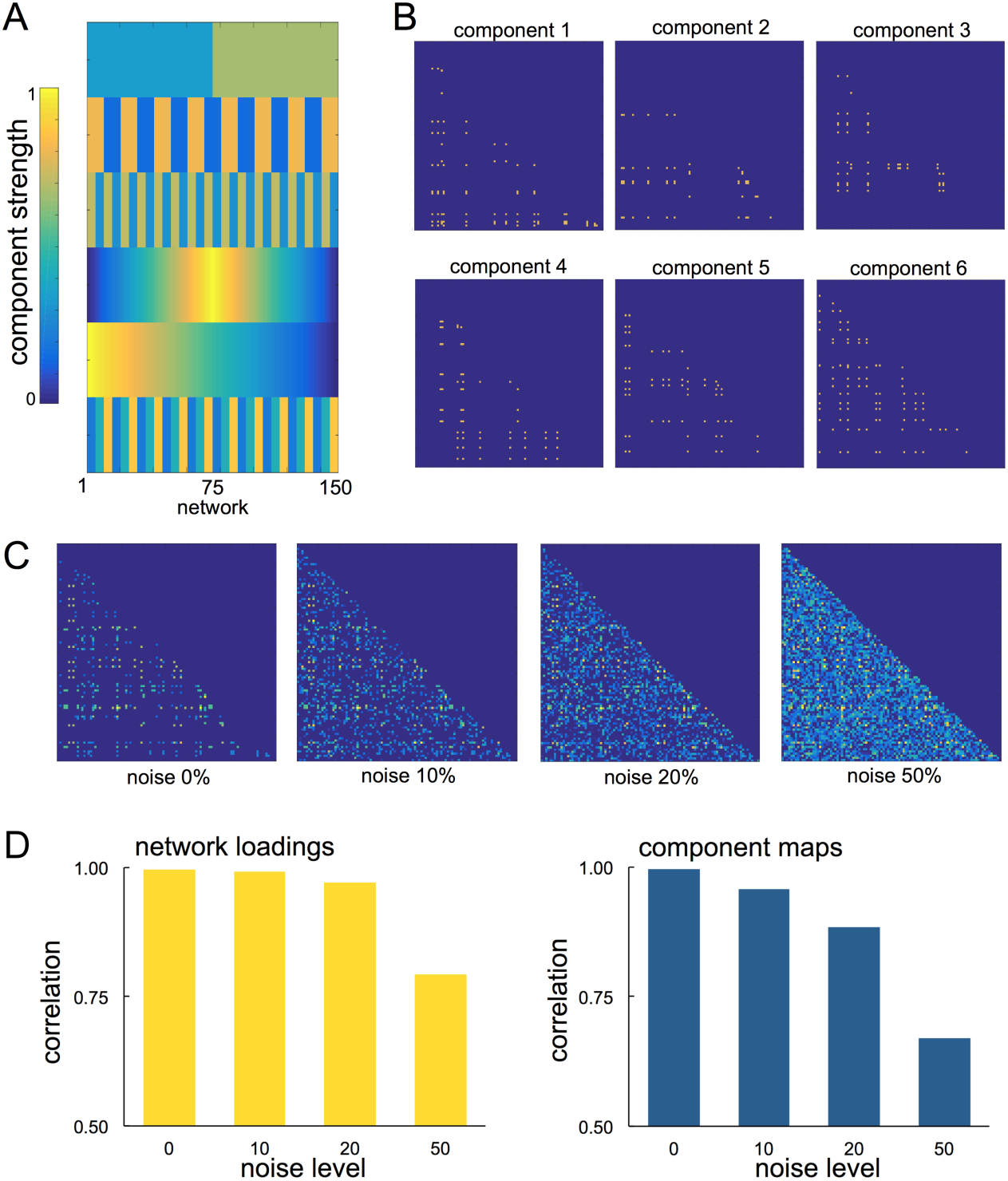
Simulating networks for PNMF decomposition. Component weights (A) and spatial maps (B) for simulating connectivity networks. Each component was weighted according to the corresponding component strength and summed to form a network. Noise was added at four levels to the final network (C). The mean correlation between recovered component loadings and the original network weights is shown in D, alongside the spatial correlation between recovered maps and the original component maps at each noise level.

For a given network, each of the six components were multiplied by the corresponding component weight and linearly summed to create the final network. Additionally, noise was added to each network by constructing a symmetric noise matrix with edge density set at 0, 10, 20 or 50%, and edge strength drawn from a normal distribution with mean and variance defined by existing network edge strengths (Figure 2C).

The simulated networks (with or without additional noise) were concatenated into data matrix V, removing any edges that were empty across all networks. ARD-PNMF was then initialised using NNSVD with rank 6 and 2500 iterations.

We found that PNMF was able to recover both the spatial maps and pattern of population variation (network loadings) of all components even under noisy conditions, achieving an average correlation between the original and recovered component maps of 0.996, 0.957, 0.884 and 0.669 (for 0, 10, 20 and 50% noise respectively) and a correlation between original and recovered component loadings of 0.997, 0.992, 0.971 and 0.794 (Figure 2D).

### Split-half reliability

To investigate if the network components can be robustly and reliability identified across population subsamples, we performed a split-half reliability assessment (Groppe et al. 2009; Groves et al. 2012). The NKI-Rockland dataset was split into two, randomly selected and equal size samples and PNMF performed independently on each. The resulting split-half components were then greedily paired with components obtained from the full dataset. Components were matched based on the correlation between the component loadings of overlapping subject populations in the half and full datasets to produce a triplet, with each original component paired with a single component from each half sample. Reliability of the original components was evaluated by measuring the edgewise (cosine) similarity between correspondent split-half component maps. Component reliability was compared to a null distribution built by randomly permuting edges in corresponding split-half pairs before calculating spatial similarity, 1000 permutations were performed for each pair.

### Rich club analysis

In order to further investigate the topological organisation of extracted network components, we performed a rich club analysis (van den Heuvel and Sporns 2011). Using the group mean structural network, nodes were sorted by degree and low degree nodes incrementally removed in steps. At each step, the density of the remaining network connections was compared to a set of 100 randomised networks of the same size to give a normalised rich club coefficient, ϕ. The rich club was defined as nodes with degree > *k*, where ϕ_*k*_ = max (ϕ). Following Collin et al., after identifying network nodes comprising the rich club, edges in each network component were defined as either ‘rich’ if they connected two rich club nodes to each other, ‘local’ if they connected two non-rich club nodes, of ‘feeder’ connections if they connected rich club nodes to non-rich club nodes (Collin et al. 2013). The overall ‘richness’ or ‘locality’ of each component was determined by comparing the number of rich or local edges to a set of 1000 equivalent random networks. Network analysis was performed with the Brain Connectivity Toolbox (Rubinov and Sporns 2010).

### Statistical analysis

For extracted components in the NKI-Rockland sample, component strength over the lifespan was modelled using polynomial regression (up to power 3) with age as a covariate and sex as an additional factor. The Akaike Information Criterion (AIC) was used to select the best model for each component (i.e.: linear, quadratic, or cubic; with or without sex). Statistical analysis was performed with the lm package in R 3.31.

In the UCLA autism cohort, component strength was compared between groups using an independent samples t-test. Analysis was performed in JASP 0.7.5.6.

### Data visualisation

The edge structure of network components were visualised with Circos (Krzywinski et al. 2009), graph nodes were ordered according to cerebral lobe and the x, y, z coordinates of the central voxel of each cerebral ROI supplied with the connectivity matrices.

In addition, to visualise the anatomical location of connected ROI in each network component, we calculated node degree (i.e.: the total number of connections of each ROI) and projected the values onto standard space masks of the cortical grey matter, basal ganglia and cerebellum. Mask voxels were assigned an ROI membership based on Euclidean distance to the nearest ROI centre and assigned the corresponding nodal degree value. Node degree images were then smoothed with a Gaussian kernel of FWHM 5mm and projected onto a 3D representation of the smoothed cortical surface using Surf Ice (www.nitrc.org/projects/surfice).

## Results

### Network components across the lifespan

#### Reliability assessment

In total, 22 network components were identified in the NKI-Rockland dataset. (Figure 4). Reliability scores were calculated for each component using a split-half framework (Groppe et al. 2009). Network components identified in two independent group samples were matched to the original set of components and the spatial similarity between the respective edge structures compared as a marker of reliability. In total, 19 components were identified in group one and 21 in group two. Of the 19 original components matched with corresponding pairs in the split-half sample, 15 demonstrated a significantly higher spatial correlation than would be expected between two random matrices with the same number of edges (all p<0.001, 1000 permutations; Table 1). The mean spatial similarity between matched components was 0.49 (range: 0.14 – 0.96). Network components identified in the full dataset are shown in Figure 3 in order of reliability.

**Table 1:** Cosine similarity between paired component maps.

| Component | Similarity | p |
| --- | --- | --- |
| A | 0.960 | 0.001 |
| B | 0.696 | 0.001 |
| C | 0.650 | 0.001 |
| D | 0.645 | 0.001 |
| E | 0.631 | 0.001 |
| F | 0.553 | 0.001 |
| G | 0.470 | 0.001 |
| H | 0.469 | 0.001 |
| I | 0.446 | 0.001 |
| J | 0.442 | 0.001 |
| K | 0.429 | 0.001 |
| L | 0.412 | 0.001 |
| M | 0.293 | 0.001 |
| N | 0.159 | 0.001 |
| O | 0.136 | 0.001 |
| P | 0.043 | 0.006 |
| Q | 0.021 | 0.228 |
| R | 0.008 | 0.671 |
| S<sup>^</sup> | - | - |
| T<sup>^</sup> | - | - |
| U<sup>^</sup> | - | - |
| V<sup>^</sup> | - | - |

<sup>^</sup> not paired with a corresponding component in

**Figure 3:**
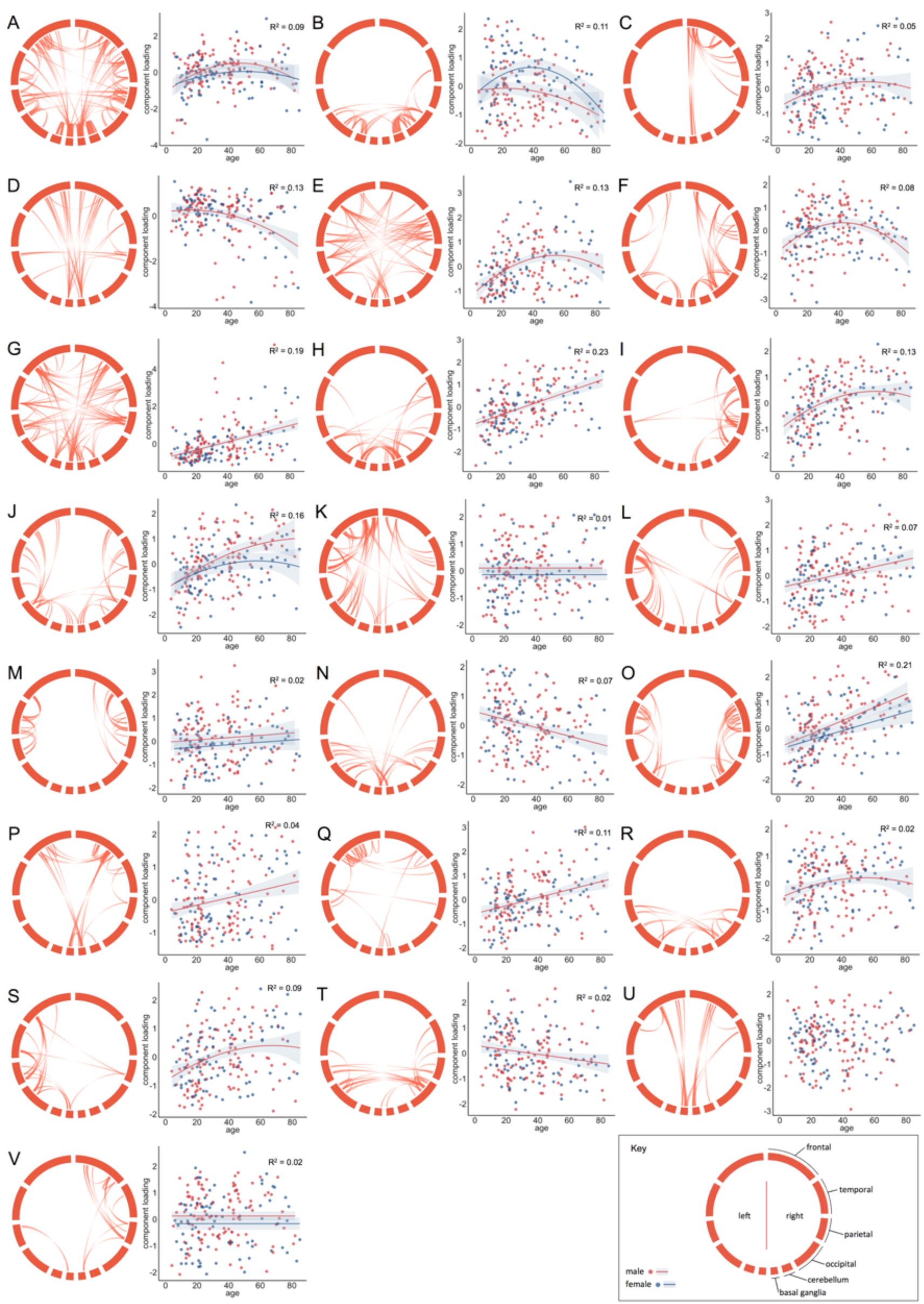
Component maps and developmental trajectories for 22 network components. PNMF decomposition in the NKI-Rockland sample resulted in 22 network components. Component maps were thresholded to retain the top 5% based on strength in the loading matrix and are displayed in circular format (see key). The relationship between (normalised) component strength and age was modelled using polynomial regression. The best model fit (see Table 2) is shown for each component. Separate model fits indicate when sex was included as an additional factor.

#### Developmental trajectories

The subject loading of each network component (i.e.: the contribution of a given component to the individual’s full network) was modelled as a function of age using polynomial regression. Of 22 components, 16 demonstrated significant age-related variation (all p<0.01; Table 2). Of these, 9 components followed nonlinear trajectories over the lifespan best described by quadratic models; 6 increased linearly with age, and 1 decreased. The addition of sex as a factor improved the model fit in 4 of the 16 significant components. Table 2 shows the best model selected for each component and Figure 4 highlights some of these trends; networks and modelled trajectories for all components are shown in Figure 3.

**Table 2:**
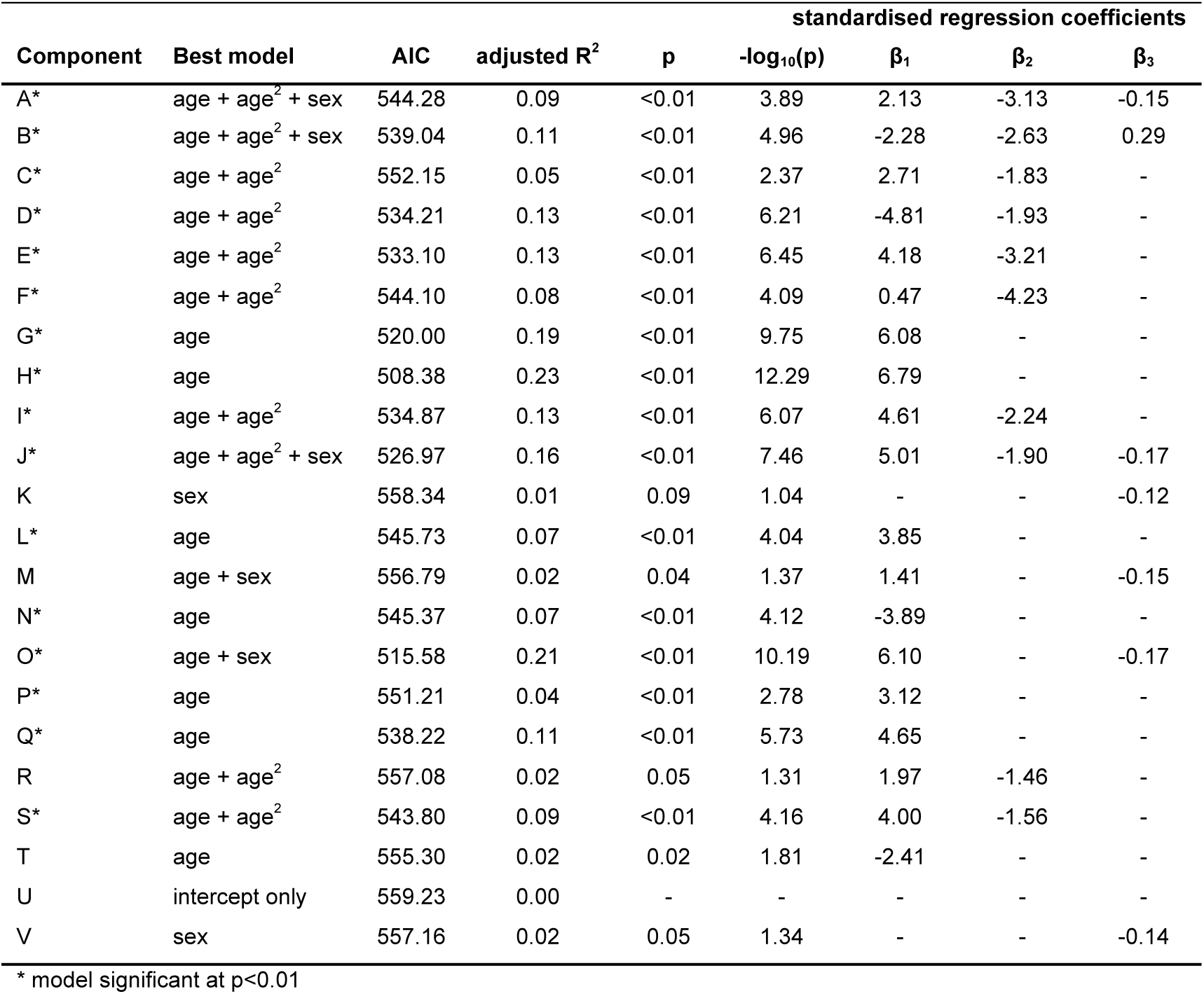
Modelling development trajectories of network components.

| Component | Best model | AIC | adjusted R<sup>2</sup> | p | standardised regression coefficients |  |  |  |
| --- | --- | --- | --- | --- | --- | --- | --- | --- |
|  |  |  |  |  | -log<sub>10</sub>(p) | β<sub>1</sub> | β<sub>2</sub> | β<sub>3</sub> |
| A* | age + age<sup>2</sup> + sex | 544.28 | 0.09 | <0.01 | 3.89 | 2.13 | -3.13 | -0.15 |
| B* | age + age<sup>2</sup> + sex | 539.04 | 0.11 | <0.01 | 4.96 | -2.28 | -2.63 | 0.29 |
| C* | age + age<sup>2</sup> | 552.15 | 0.05 | <0.01 | 2.37 | 2.71 | -1.83 | - |
| D* | age + age<sup>2</sup> | 534.21 | 0.13 | <0.01 | 6.21 | -4.81 | -1.93 | - |
| E* | age + age<sup>2</sup> | 533.10 | 0.13 | <0.01 | 6.45 | 4.18 | -3.21 | - |
| F* | age + age<sup>2</sup> | 544.10 | 0.08 | <0.01 | 4.09 | 0.47 | -4.23 | - |
| G* | age | 520.00 | 0.19 | <0.01 | 9.75 | 6.08 | - | - |
| H* | age | 508.38 | 0.23 | <0.01 | 12.29 | 6.79 | - | - |
| I* | age + age<sup>2</sup> | 534.87 | 0.13 | <0.01 | 6.07 | 4.61 | -2.24 | - |
| J* | age + age<sup>2</sup> + sex | 526.97 | 0.16 | <0.01 | 7.46 | 5.01 | -1.90 | -0.17 |
| K | sex | 558.34 | 0.01 | 0.09 | 1.04 | - | - | -0.12 |
| L* | age | 545.73 | 0.07 | <0.01 | 4.04 | 3.85 | - | - |
| M | age + sex | 556.79 | 0.02 | 0.04 | 1.37 | 1.41 | - | -0.15 |
| N* | age | 545.37 | 0.07 | <0.01 | 4.12 | -3.89 | - | - |
| O* | age + sex | 515.58 | 0.21 | <0.01 | 10.19 | 6.10 | - | -0.17 |
| P* | age | 551.21 | 0.04 | <0.01 | 2.78 | 3.12 | - | - |
| Q* | age | 538.22 | 0.11 | <0.01 | 5.73 | 4.65 | - | - |
| R | age + age<sup>2</sup> | 557.08 | 0.02 | 0.05 | 1.31 | 1.97 | -1.46 | - |
| S* | age + age<sup>2</sup> | 543.80 | 0.09 | <0.01 | 4.16 | 4.00 | -1.56 | - |
| T | age | 555.30 | 0.02 | 0.02 | 1.81 | -2.41 | - | - |
| U | intercept only | 559.23 | 0.00 | - | - | - | - | - |
| V | sex | 557.16 | 0.02 | 0.05 | 1.34 | - | - | -0.14 |

\* model significant at p<0.01

**Figure 4:**
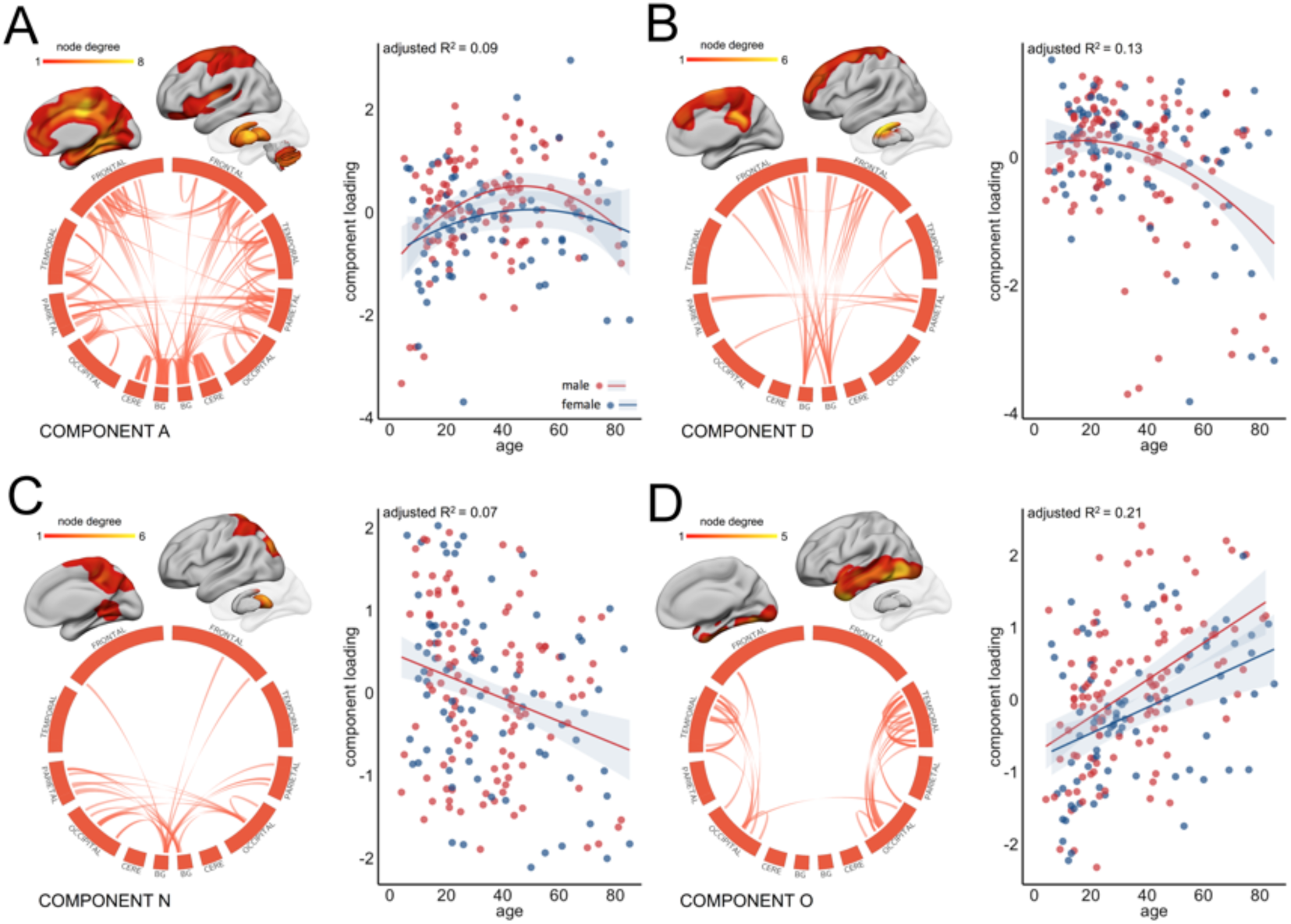
Age-related variation in component strength. Four network components are highlighted as examples of age-related variation in subnetwork connectivity (see Fig 4 for all components). For visualisation, components maps were thresholded at the 95^th^ percentile and connections are shown in circular format. To show the anatomical location of connected regions, node degree was calculated for each cerebral ROI as the sum of its connections in the thresholded component and projected onto cortical/subcortical surfaces. The relationship between (normalised) component strength and age was modelled using polynomial regression. The best model fit is shown for each component. Separate model fits indicate when sex was included as an additional factor. Red indicates male; blue, female. Cere=cerebellum, BG=basal ganglia.

In Figure 4, component A comprises a robust (split-half similarity: 0.96) and relatively dense pattern of connectivity including both local, within-lobe, and longer, between-lobe, connections. Interhemispheric connections are apparent between frontal and parietal lobes. Highly connected cerebral regions in this component include the cingulate and paracingulate, insular, medial temporal, and superior parietal cortices, with dense connectivity also evident within the basal ganglia, and between the basal ganglia and higher cerebral cortex. Over the lifespan, connectivity between these regions increases rapidly over childhood and adolescence, peaking between 40 and 50 and declining into older age. Component loading is slightly higher in males than in females across the lifespan though follows a steeper decline with old age. Similar trends in connectivity are seen in components E, F and J (Fig 3). Components D and N (Fig 4B; 4C) capture bilateral patterns of connectivity between subcortical grey matter structures and frontal and parietal cortices, respectively. Edges in component D predominantly connect the caudate nucleus and superior, medial frontal regions. The strength of this component remains relatively stable until middle age before declining. The strength of component N (Fig 4C) monotonically decreases with age with edges connecting the thalamus to post-central cortex and superior parietal regions. In contrast, component O follows a linearly increasing trend with age (Fig 4D) Connections in this component are primarily local, connecting anatomically adjacent cortical regions within the temporal lobe and temporo-occipital junction in both hemispheres. A similar pattern can be seen in component L (Fig 3).

#### Component topology

Qualitative assessment of the patterns of connectivity within components reveals a generally bilateral and symmetric organisation across hemispheres. Unilateral components appear to reflect the topology of corresponding components in the opposite hemisphere (e.g; I and S; O and L).

In order to quantify whether inter-regional connections within individual network components revealed a preferential support for connectivity between network hubs over topologically local nodes, we performed a rich club analysis. Sixty-five nodes with degree greater than 100 were defined as the rich club (maximum ϕ = 1.09) and included nodes bilaterally in: frontal pole; cingulate and paracingulate cortex; insula; hippocampus; precuneus and superior parietal lobule; lateral occipital cortex, and basal ganglia structures (caudate, pallidum and thalamus).

Component edges were defined as rich, feeder or local connections, and the ‘richness’ or ‘locality’ of each component defined as the number of rich or local connections compared to a set of 1000 random networks (Table 3). Four components were found to contain significantly more rich-club connections than expected by chance (A, E, F and I; all p=0.001). These components are shown in Figure 5; richness and locality indices for all components are shown in Table 3. All rich components show a similar inverted ‘U’ trajectory with age (Table 1, Figure 3). Edge strength distributions in Figure 5 show that component richness is associated with both an increased number and strength of rich compared to local edges. In contrast, four components were to found to have significantly more local connections than expected by chance (Fig 5; M, O, L and J; p=0.001). These components all had increased number and strength of local compared to rich connections and increased in strength across the lifespan (Table 1, Figure 3).

**Table 3:** Rich club analysis of network components.

| Component | Number of edges | Rich edges | Richness<sup>^</sup> | p | Local edges | Locality<sup>^</sup> | p |
| --- | --- | --- | --- | --- | --- | --- | --- |
| A | 2549 | 780 | 1.34 | 0.001 * | 658 | 1.00 | 0.418 |
| B | 788 | 115 | 0.64 | 1.000 | 223 | 1.10 | 0.035 |
| C | 530 | 61 | 0.50 | 1.000 | 152 | 1.12 | 0.059 |
| D | 605 | 155 | 1.12 | 0.049 | 134 | 0.87 | 0.987 |
| E | 1119 | 434 | 1.69 | 0.001 * | 158 | 0.55 | 1.000 |
| F | 835 | 238 | 1.25 | 0.001 * | 169 | 0.79 | 1.000 |
| G | 1583 | 396 | 1.09 | 0.015 | 333 | 0.82 | 1.000 |
| H | 757 | 160 | 0.92 | 0.882 | 211 | 1.08 | 0.071 |
| I | 520 | 155 | 1.30 | 0.001 * | 119 | 0.89 | 0.917 |
| J | 623 | 73 | 0.51 | 1.000 | 193 | 1.21 | 0.001 * |
| K | 1400 | 293 | 0.91 | 0.967 | 309 | 0.86 | 1.000 |
| L | 628 | 86 | 0.60 | 1.000 | 210 | 1.30 | 0.001 * |
| M | 576 | 44 | 0.33 | 1.000 | 254 | 1.71 | 0.001 * |
| N | 535 | 119 | 0.97 | 0.612 | 103 | 0.75 | 1.000 |
| O | 761 | 93 | 0.53 | 1.000 | 316 | 1.62 | 0.001 * |
| P | 710 | 161 | 0.99 | 0.540 | 184 | 1.01 | 0.446 |
| Q | 498 | 111 | 0.98 | 0.599 | 153 | 1.19 | 0.007 |
| R | 342 | 70 | 0.89 | 0.830 | 94 | 1.07 | 0.236 |
| S | 572 | 132 | 1.01 | 0.425 | 120 | 0.82 | 0.999 |
| T | 599 | 157 | 1.15 | 0.019 | 132 | 0.86 | 0.987 |
| U | 481 | 41 | 0.37 | 1.000 | 120 | 0.97 | 0.648 |
| V | 345 | 55 | 0.70 | 0.999 | 100 | 1.13 | 0.073 |

<sup>^</sup> number of edges compared to 1000 random networks of the same size

**Figure 5:**
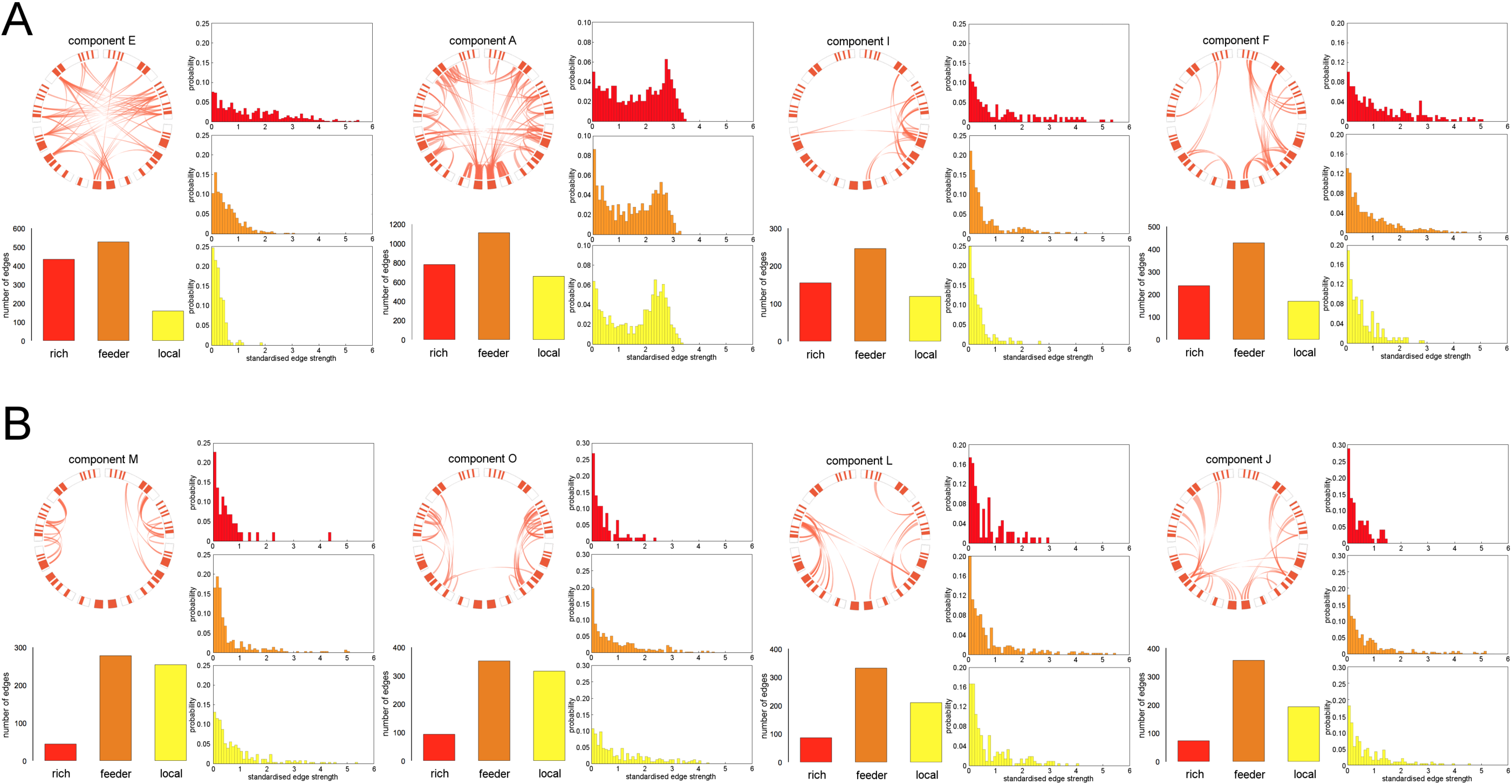
Rich club analysis of network components. Network components with significantly more rich-club (A) or local (B) edges than in a set of 1000 equivalent random networks are shown. Thresholded connectivity maps are displayed in circular diagrams as in Fig 4, with rich club nodes highlighted in red. The number and probability distribution of rich, feeder and local edges are displayed for each component.

### Network components in autism spectrum disorder

In our second experiment, we applied PNMF to a dataset comprising high-functioning individuals with Autism Spectrum Disorder and a set of age- and sex-matched controls. In total, 24 components were identified (Table 4). One component was found to have a significantly higher loading in the ASD group compared to typically developing individuals (t_(1,92)_=3.161; p=0.002; p<0.05 after Bonferroni correction for multiple comparisons). Additionally controlling for age did not alter this relationship (ANCOVA: F_(1,91)_ = 11.05, p<0.001). To control for possible gender effects, we repeated the statistical analysis after excluding female participants (n=7 TD; n=6 ASD). The difference in component strength remained significant (t_(1_,_79)_=344; p<0.001).

**Table 4:** Between-group comparison of component loadings.

| Component | $t_{(1,92)}$ | p | Cohen's $d$ |
| --- | --- | --- | --- |
| 1 | 1.769 | 0.080 | 0.366 |
| 2 | 1.576 | 0.119 | 0.326 |
| 3 | 0.809 | 0.421 | 0.168 |
| 4 | 0.308 | 0.759 | 0.064 |
| 5 | 1.388 | 0.168 | 0.287 |
| 6 | -0.735 | 0.464 | -0.152 |
| 7 | -0.342 | 0.733 | -0.071 |
| 8 | -1.180 | 0.241 | -0.244 |
| 9 | 0.698 | 0.487 | 0.145 |
| 10 | 3.161 | 0.002* | 0.654 |
| 11 | 0.730 | 0.467 | 0.151 |
| 12 | 1.421 | 0.159 | 0.294 |
| 13 | -0.095 | 0.925 | -0.020 |
| 14 | -0.019 | 0.985 | -0.004 |
| 15 | 1.670 | 0.098 | 0.346 |
| 16 | -0.376 | 0.708 | -0.078 |
| 17 | 0.146 | 0.884 | 0.030 |
| 18 | 0.328 | 0.744 | 0.068 |
| 19 | 0.283 | 0.778 | 0.059 |
| 20 | 0.162 | 0.872 | 0.033 |
| 21 | 1.459 | 0.148 | 0.302 |
| 22 | 1.162 | 0.248 | 0.241 |
| 23 | 0.410 | 0.683 | 0.085 |
| 24 | -0.528 | 0.599 | -0.109 |

\* significant at $p<0.05$ after Bonferroni correction for multiple comparisons

This component is shown in detail in Figure 6 and comprised a bilateral and symmetric pattern of connectivity with edges predominantly linking nodes in the anterior and posterior cingulate cortex, paracingulate cortex, supplemental motor areas, and parietal cortex. Both intra- and interhemispheric connections are visible with additional connections between the putamen and parietal cortex in both hemispheres. The mean (±S.D.) component loadings were 2.9 ± 0.69 in the ASD group and 2.5 ± 0.57 in the typically-developing group (Fig 6B). Rich club analysis revealed that this component had significantly more local connections than expected by chance (locality: 1.32, p=0.001).

**Figure 6:**
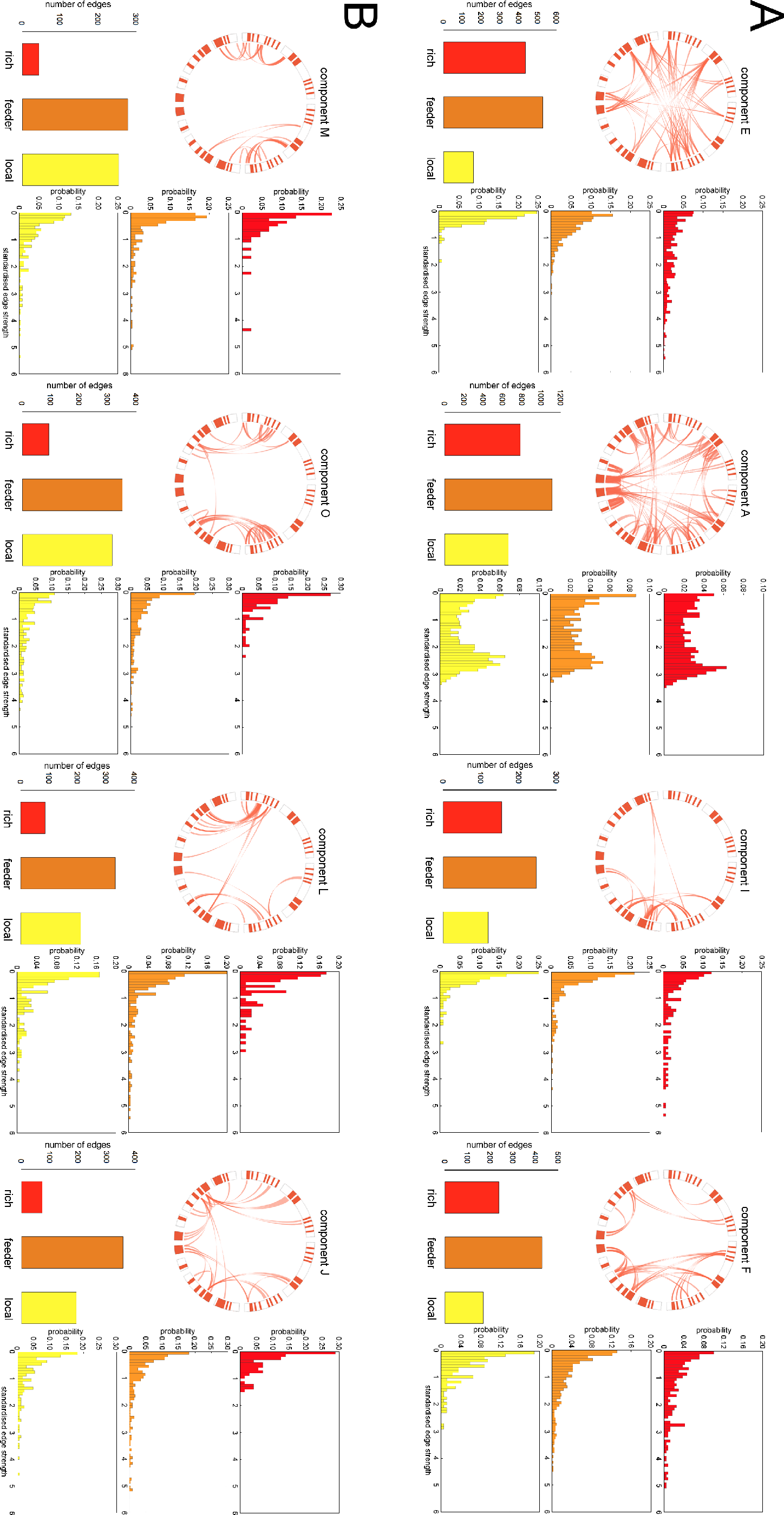
Structural connectivity is significantly increased in ASD. A single subnetwork was found to be significantly stronger in the ASD cohort. The component map is shown in A (thresholded at 95^th^ percentile), and component loadings for both groups compared in B. The anatomical locations of connected regions are visualised as in Fig 6 and shown in C.

As noted above, a threshold was applied to the network data before performing PNMF, limiting the analysis to edges shared by at least 10% of participants. We additionally performed PNMF after applying two alternate thresholds: 5% and 20%. At both thresholds, a similar pattern was observed and found to be significantly different between groups (at 5%: t=2.732, p=0.008; excluding females: t=3.16, p=0.002; at 20%: t=2.40, p=0.018; excluding females: t=2.77, p=0.007). The spatial patterns associated with this component at each threshold match closely to that described above.

## Discussion

In this study, we model structural connectivity networks as the combination of separable subnetworks, or network components, using a data-driven and multivariate approach. We show that network components can be reliably identified across individuals, follow a developmental trajectory with age and highlight differences in connectivity in autism spectrum disorder.

NMF provides a natural setting for the analysis of neuroimaging data due to the inherent nonnegativity common to many imaging-derived metrics (e.g.: tissue volume, fibre count) (Sotiras et al. 2015). In image analysis, NMF leads to a parts-based representation of the data, extracting sparse image components with localised spatial support (Lee and Seung 1999). In this way, NMF confers a relatively simple interpretation of the data, namely that the complex whole can be approximated by the summation of the localised parts. We show that decomposing structural connectivity networks with NMF results in a soft clustering of connections that co-vary together across the population forming relatively sparse subnetworks. We also find that these components are biologically relevant, capturing known subsystems (e.g.: subcortical-cortical projections; components C, D, N and P), local patterns of connectivity between anatomically or functionally homologous regions (e.g.: components L, O and T) and varying with age across the lifespan.

MR studies have found that both white and grey matter tissue volumes follow an inverted U trajectory over the human lifespan, rising rapidly in development, peaking in the second or third decade and declining into older age (Giedd et al. 1999; Westlye et al. 2010; Ziegler et al. 2012). Markers of microstructural maturation in cerebral white matter also follow similar trends (Westlye et al. 2010; Lebel et al. 2012). In the NKI-Rockland cohort, a recent graph theoretical analysis of structural connectivity found several network properties, including efficiency, varied along the similar polynomial trajectories (Zhao et al. 2015). In contrast, the modularity – or extent to which a network could be segregated into regional communities – remained stable over time.

In this study, using a subsample of the NKI-Rockland dataset, we found that several network components followed an inverted U-shaped trajectory. Significant variation in component strength with age was found in 16 of 22 components, of which 9 followed a nonlinear trajectory. When ordered by reliability across subsamples, age variation in 8 of the top 10 most robust components was best described by a polynomial relationship. Of the remaining components, 6 increased linearly with age, and 1 decreased. We found that components linking densely-connected network hub regions tended to follow a nonlinear trajectory with age, decreasing in strength in later life. In contrast, components that increased in strength with age reflected local connectivity patterns between neighbouring regions, or between corresponding regions in the opposite hemisphere (e.g. components H, L, M, O and Q), compared to more global connectivity patterns (A, E, F, J).

These observations support evidence of an increasing dependence upon local connectivity in the elderly connectome with corresponding decreases in long-range connection strength between cortical hubs (Betzel et al. 2014; Cao et al. 2014; Perry et al. 2015; Zhao et al. 2015). Analyses of functional connectivity networks have shown that hub connectivity follows a U shaped trajectory over the human lifespan alongside a decrease in network modularity, suggesting a less segregated network topology in old age (Cao et al. 2014; Chan et al. 2014). Performing a comparative analysis of both functional and structural networks, Betzel et al. showed structural connectivity of hub regions decreases dramatically with age, while local connectivity is relatively spared. In addition, functional connectivity within resting state networks (RSN) decreased with age whereas connectivity between RSN increased. This increase in between-network connectivity was subserved by multi-step structural connections. Similarly,Perry et al showed that alterations to edge strength in the aging structural connectome likely lead to an increased preference for network communication via multiple, non-hub, local pathways (Perry et al. 2015). Taken together, this evidence suggests that the decline of long-distance, hub-to-hub connections with a relative sparing of topologically, local connections results in less efficient network communication in older age, a process that a may underlie progressive cognitive decline (O’Sullivan et al. 2001; Andrews-Hanna et al. 2007)

In our second experiment, we found a single subnetwork with significantly greater connectivity on average in an adolescent ASD cohort. This network was composed primarily of multiple, local connections between neighbouring regions in the cingulate and paracingulate cortices, both within and between hemispheres, along with connections between supplementary motor areas, parietal cortex and the putamen. The cingulum, and in particular the anterior cingulate cortex, has been linked to ASD due mainly to its role in social interaction and attention (Mundy 2003). Similarly, metabolic (Tebartz van Elst et al. 2014), neuropathological (Simms et al. 2009) and neuroanatomical disturbances (Schumann et al. 2010) have also been reported in the cingulate in ASD. In a comprehensive diffusion tractography study of the white matter tracts of the limbic system, Pugliese et al. found significantly increased tract volume bilaterally in the cingulum bundle (determined by streamline count) of adults with Asperger’s syndrome (Pugliese et al. 2009). This observation is convergent with post-mortem findings in autistic cases of excessive axonal connectivity between neighbouring cortical regions in the cingulate (Zikopoulos and Barbas 2010). This is of particular interest, given the putative neurodevelopmental origins of ASD, as the cingulum bundle forms early in gestation, followed by short range cortico-cortical connections in the third trimester (Vasung et al. 2010; Takahashi et al. 2012) suggesting that any early disruptions to white matter development in utero could have significant long-term implications on neurodevelopment.

Previous structural network analyses in autistic populations have found significantly increased structural connectivity, inferred from tractography streamline count, amongst cortical regions including anterior and posterior cingulate cortex, superior frontal, superior parietal, and insula cortex (Ray et al. 2014). In a subset of the UCLA cohort, Watanabe and Rees found evidence for delayed, or immature, hub connectivity in ASD (Watanabe and Rees 2015), whereas Ghanbari et al. used a supervised variant of NMF coupled with a graph embedding approach to define discriminative network components and found decreased interhemispheric connection strength in subcortical subnetworks in ASD (Ghanbari et al. 2014). Similarly, other studies have reported decreases in structural connectivity, dependent on measures of white matter microstructure (Thakkar et al. 2008; Noriuchi et al. 2010; Lo et al. 2011). These discrepancies may relate, in part, to the uncertain correspondence between different measures of structural connectivity. In the original study of this cohort, Rudie et al. previously found streamline count was significantly increased in 4 times as many connections in ASD subjects compared to controls (Rudie et al. 2012). However, a concomitant decrease in fractional anisotropy and increase in mean diffusivity was also noted in white matter connections on average in ASD. These differences resulted in an atypical age-related development of network efficiency in ASD, a factor that related to symptom severity (Rudie et al. 2012).

Non-negative matrix factorisation belongs to a class of multivariate matrix decomposition and dimension reduction techniques that include principal component analysis (PCA) and independent component analysis (ICA). Recently, exploratory multivariate analysis methods have proven well-suited to the discovery of complex organisational relationships in the brain (Beckmann and Smith 2005; Calhoun et al. 2009; McIntosh and Mišić 2013). Previous studies have shown the potential of matrix factorisation techniques to isolate topological subnetworks from functional and structural connectivity matrices on an individual or group level (Clayden et al. 2013; Ghanbari et al. 2014; Park et al. 2014). We performed simulations to demonstrate that PNMF is particularly able to retrieve superposed spatial patterns and the corresponding, population-varying component weights from a set of noisy connectivity networks.

Additionally, an important aspect of any dimension reduction task is determining the optimal dimensionality of the solution. In this paper, we chose to employ an automatic model selection process that iteratively updates model rank, removing components with low spatial variance that do not contribute significantly to the final reconstruction (Yang et al. 2010). This resulted in around 20 effective components selected in both experimental cohorts. Previous studies have shown that, for ICA-based decomposition of functional MRI data, higher-dimensional decompositions can reveal nested subsystems within functional networks (Kiviniemi et al. 2009; Abou Elseoud et al. 2011; Dipasquale et al. 2015). Indeed, in the present study we observed apparently correspondent components that were split between opposite hemispheres (e.g.: components I and S), whereas others formed bilateral symmetric patterns (components A, U and H) suggesting some hierarchical structure within components. PNMF network decomposition at multiple dimensionalities would provide a framework to investigate the nested, or hierarchical nature of structural connectivity subnetworks (Meunier et al. 2010; Betzel et al. 2013; Betzel and Bassett 2016).

### Study limitations

The use of (log-transformed) streamline counts to estimate structural connectivity could be considered a limitation of this study. Is it important to note that fibre counts do not necessarily reflect true anatomical connectivity, and tractography is prone to mapping false positive connections due to local accumulation of modelling errors (Jones 2010; Jones et al. 2013). However, using available open access datasets, we have shown that PNMF is able to extract subnetworks that can provide insight into biological variability, with developmental trajectories that suggest streamline count, in part, can reflect maturational processes in the brain. Importantly, PNMF is generalisable to any inherently non-negative data. This opens future avenues to explore network components derived from modern, probabilistic tractographic algorithms that better reflect true anatomical connectivity (Pestilli et al. 2014; Smith et al. 2015).

### Conclusions

In conclusion, we present a multivariate analysis of structural connectivity in two cohorts. We demonstrate that complex networks can be decomposed into robust and reliable subnetworks that vary in strength with age. Further, we identify a specific subnetwork with increased connection strength in autism spectrum disorder. We propose that this form of network component analysis shows good potential for further exploration of the human structural connectome.

## Acknowledgements

The authors would like to thank Dr Jesse Brown and all involved in the collection and aggregation of data for the USC Multimodal Connectivity Database. This research was conducted within the Developmental Imaging research group, Murdoch Childrens Research Institute and the Children’s MRI Centre, Royal Childrens Hospital, Melbourne, Victoria. It was supported by the Murdoch Childrens Research Institute, the Royal Children’s Hospital, Department of Paediatrics The University of Melbourne and the Victorian Government s Operational Infrastructure Support Program. The project was generously supported by RCH1000, a unique arm of The Royal Children’s Hospital Foundation devoted to raising funds for research at The Royal Children’s Hospital.

